# Temporal Dynamics of Azot Expression Following Traumatic Brain Injury

**DOI:** 10.1101/2025.04.29.651334

**Authors:** Andrés Gutiérrez-García, Mariana Marques-Reis, Barbara Hauert, Eduardo Moreno

## Abstract

Cell competition, a conserved biological process in which cells compete for survival based on relative fitness, has emerged as a critical mechanism in diverse biological contexts. Here, we investigate the role of cell competition in traumatic brain injury (TBI) by characterizing the temporal expression pattern of Azot, a key downstream effector of fitness-based selection, following injury in *Drosophila melanogaster*. Our findings reveal a distinct temporal profile of Azot expression post-TBI, with TUNEL assays confirming that Azot-expressing cells undergo apoptotic elimination. We demonstrate that following injury, the proportion of dying cells marked as “losers” significantly increases compared to non-injured conditions, indicating that cell competition becomes a predominant elimination mechanism during acute post-injury phases. Contrary to previous findings in neurodegenerative disease models where competition was restricted to neurons, we show that following TBI, both neurons and glia are subject to competitive elimination. Furthermore, in *azot* knockout conditions, we observe an accumulation of cells attempting to express Azot, suggesting impaired clearance of suboptimal cells. These findings advance our understanding of cellular quality control mechanisms following brain injury and may inform the development of novel therapeutic approaches to enhance functional recovery after TBI.

## Introduction

Cell competition is a conserved biological process in which cells within a tissue compete for survival based on their relative fitness. First described by Morata and Ripoll in 1975, this phenomenon was initially observed in *Drosophila melanogaster*, where slower-growing *minute* mutant cells were eliminated from developing wing disc tissue when juxtaposed with wild-type cells1. The significance of cell competition remained underappreciated until 2002, when Moreno and colleagues revealed its potential tumor-suppressive function by demonstrating how it facilitates the detection and elimination of aberrant cells^2^. They established that, in *minute* mutants, reduced access to Decapentaplegic (Dpp) morphogen leads to upregulation of the transcriptional repressor *brinker*, ultimately triggering c-Jun N-terminal kinase (JNK)-dependent apoptosis^2^.

Subsequent research has expanded our understanding of cell competition across diverse biological contexts, including development^3,4^, tissue homeostasis^4–6^, aging^7–10^, and cancer^11– 14^. Three principal mechanisms underlie this process: competition for survival factors, competition through fitness fingerprints, and mechanical cell competition^14–16^. The identification of Flower proteins as “fitness fingerprints” was particularly significant, establishing the molecular basis for how cells communicate their fitness status to neighboring cells^17,18^.

In *Drosophila*, the *flower* gene produces three protein isoforms: Flower Ubi, which marks winner cells, and Flower LoseA and Flower LoseB, which typically mark loser cells— although Flower LoseA functions as a winner marker in the brain^18,19^. The human genome encodes four Flower proteins (two winner, two loser) that serve as prognostic markers in various pathologies, including cancer progression and COVID-19 severity^20,21^.

A critical downstream effector in fitness fingerprint-mediated cell competition is the protein Azot, which functions as a fitness checkpoint downstream of Flower and upstream of the pro-apoptotic gene *hid*^8^. Studies have demonstrated the significance of Azot in multiple physiological and pathological processes, particularly in aging^8,22^ and neurodegeneration^7^, where cell competition in the brain was circumscribed to occurring only in neurons^7^. Cells expressing Flower LoseB activate Azot, which subsequently triggers apoptotic pathways, resulting in the elimination of viable yet suboptimal cells from the tissue. Recent literature challenges the idea that the lack of *azot* blocks cell competition^10^; instead, the accumulation of loser cells in the absence of *azot* is likely the result of a slower elimination mechanism rather than complete inhibition of the competitive process^10^.

Traumatic brain injury (TBI) represents a major global health challenge, characterized by brain damage resulting from external forces^23^. The pathophysiology of TBI involves two distinct phases: primary injury, occurring at the moment of impact with direct damage to neural tissue and vasculature; and secondary injury, encompassing cascades of molecular and cellular events including inflammation, oxidative stress, and cellular dysfunction that can persist for days following the initial trauma^24^.

Recent studies in *Drosophila* have demonstrated that TBI induces upregulation of Flower LoseB in specific cells within the injured brain region of adult flies^25–27^. This observation proves that selection mechanisms influence cellular fate following brain damage. Furthermore, Azot expression has been detected in these loser cells following TBI^27^, raising important questions about the temporal dynamics of this expression and the consequences of disrupting the cell competition process through *azot* depletion.

The relationship between cell competition and TBI recovery represents an understudied yet potentially crucial aspect of post-injury tissue remodeling. While inflammation and immune responses following TBI have been extensively characterized^23,28^, the role of fitness-based selection mechanisms in determining which cells survive or die remains poorly understood. This knowledge gap is particularly significant given emerging evidence that selective elimination of suboptimal cells may be essential for optimal tissue function and recovery^7^.

Understanding how Azot-mediated elimination of loser cells contributes to post-TBI tissue homeostasis could provide insights into novel therapeutic approaches. If cell competition serves a beneficial role in clearing damaged or dysfunctional cells, enhancing this process might improve recovery outcomes^7^. Conversely, if certain aspects of cell competition exacerbate secondary injury, targeted inhibition might preserve salvageable tissue.

In this study, we characterize the temporal profile of Azot expression following TBI and investigate the consequences of disrupting cell competition through *azot* depletion. Additionally, we examine which cell populations are subject to competition-based elimination in the injured brain. Our findings provide new insights into the cellular dynamics of post-injury brain tissue and suggest potential avenues for therapeutic intervention in TBI.

## Results

### Azot is expressed in the wounded area upon TBI, leading to loser cell elimination

Knowing that Flower LoseB and Azot are expressed upon TBI^26,27^, we wanted to investigate the temporal expression of Azot following injury. To examine this, we performed a stab lesion procedure with a filament next to the pair of bristles surrounding the retina^25,26^. This method is preferable to injuring through the retina, as the filament can carry remnants of damaged photoreceptors inward, causing undesirable autofluorescence, particularly in the “red” spectrum, due to the presence of metarhodopsin^29,30^.

We assessed Azot expression at 24, 48, 72 hours, and 7 days after damage (AD) and compared it to a non-injured control of similar age (Figure 1). Our results demonstrate that the number of Azot-positive cells peaks within the first 24 hours following TBI (mean of 80.2 cells per optic lobe [OL]), after which it gradually decreases over time. At 48 hours AD, there was a 12.5% reduction in the number of Azot-expressing cells (mean of 70.67 cells per OL) compared to 24 hours AD, and at 72 hours AD, a further 21.5% decrease (mean of 55.3 cells per OL) compared to 48 hours AD. By the 7th day after injury, levels of Azot-positive cells (mean of 16 cells per OL) were no longer significantly different from those observed in a non-injured scenario (mean of 9.5 cells per OL).

**Figure 1.**
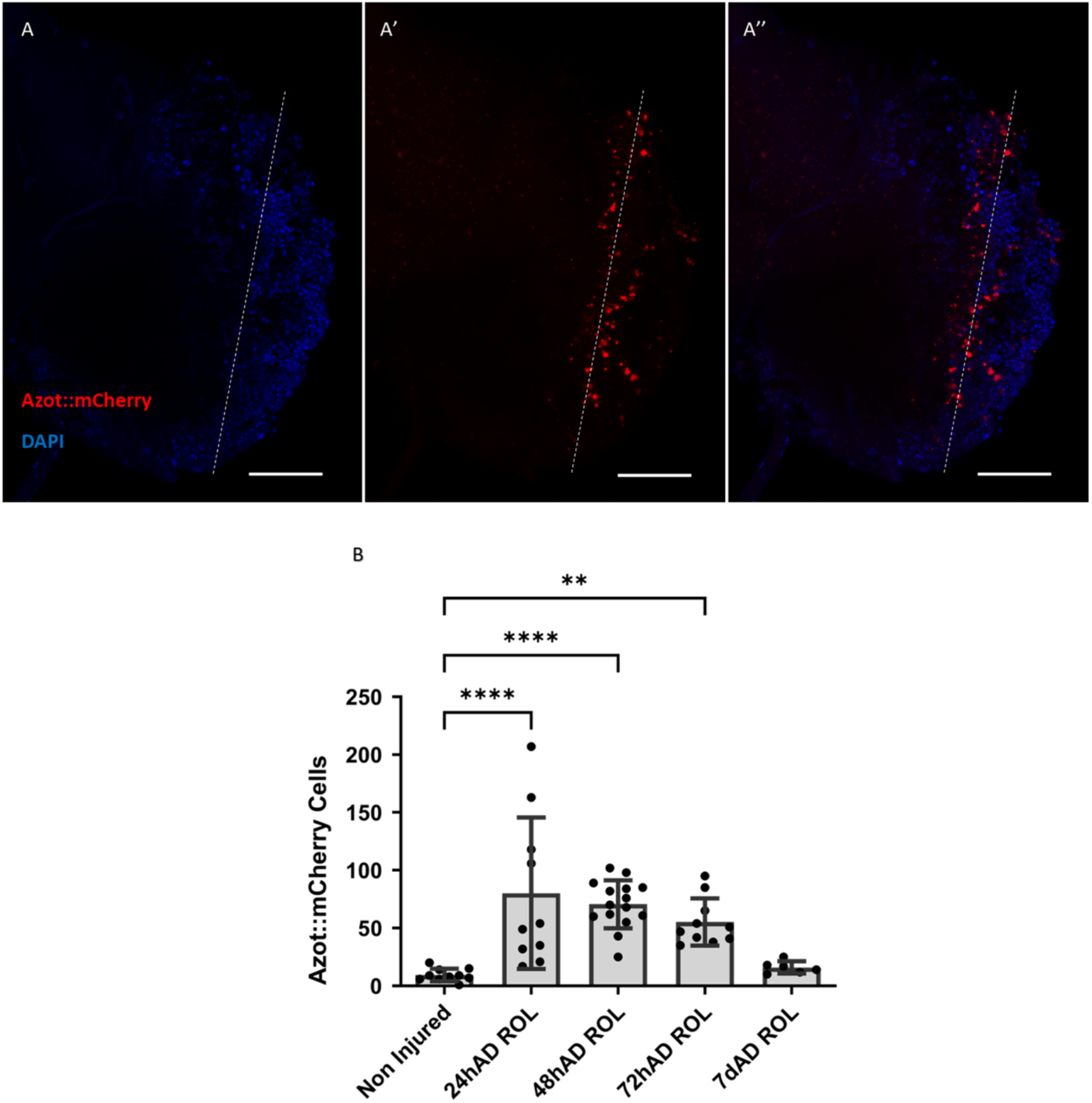
Temporal expression of Azot upon TBI. (A) Inset of a ROL 48h after TBI. A is the blue channel for DAPI, A’ is the red channel for mCherry, and A” is the merge of blue and red channels. The dashed line indicates the wounded area. 40µm Maximum intensity projection. Scale bar: 50µm. (B) Quantification of the number of Azot::mCherry expressing cells in the OL. Statistical comparisons shown are only made with the “Non-injured” control. Error bars indicate SD; **P≤0.01; ****P≤0.0001. Statistical significance was calculated with one-way ANOVA and Dunnett’s post hoc test. Genetic background: *ywf*; *azot*{KO-KI *azot::mCherry*} (Hz); +

We then addressed cell death after TBI using a TUNEL assay to evaluate cell death at 24, 48, and 72 hours AD (Figure 2A). Our results reveal that TUNEL-positive cells increase over time after damage. In the first 24 hours following TBI, there was a 449.4% increase in TUNEL-positive cells (mean of 14.67 cells per OL) compared to the control (mean of 2.67 cells per OL). At 48 hours AD, there was a 72.7% higher number of TUNEL-positive cells (mean of 25.33 cells per OL) than at 24 hours AD, and at 72 hours AD, approximately 13.8% more TUNEL-positive cells (mean of 28.83 cells per OL) than at 48 hours AD.

**Figure 2.**
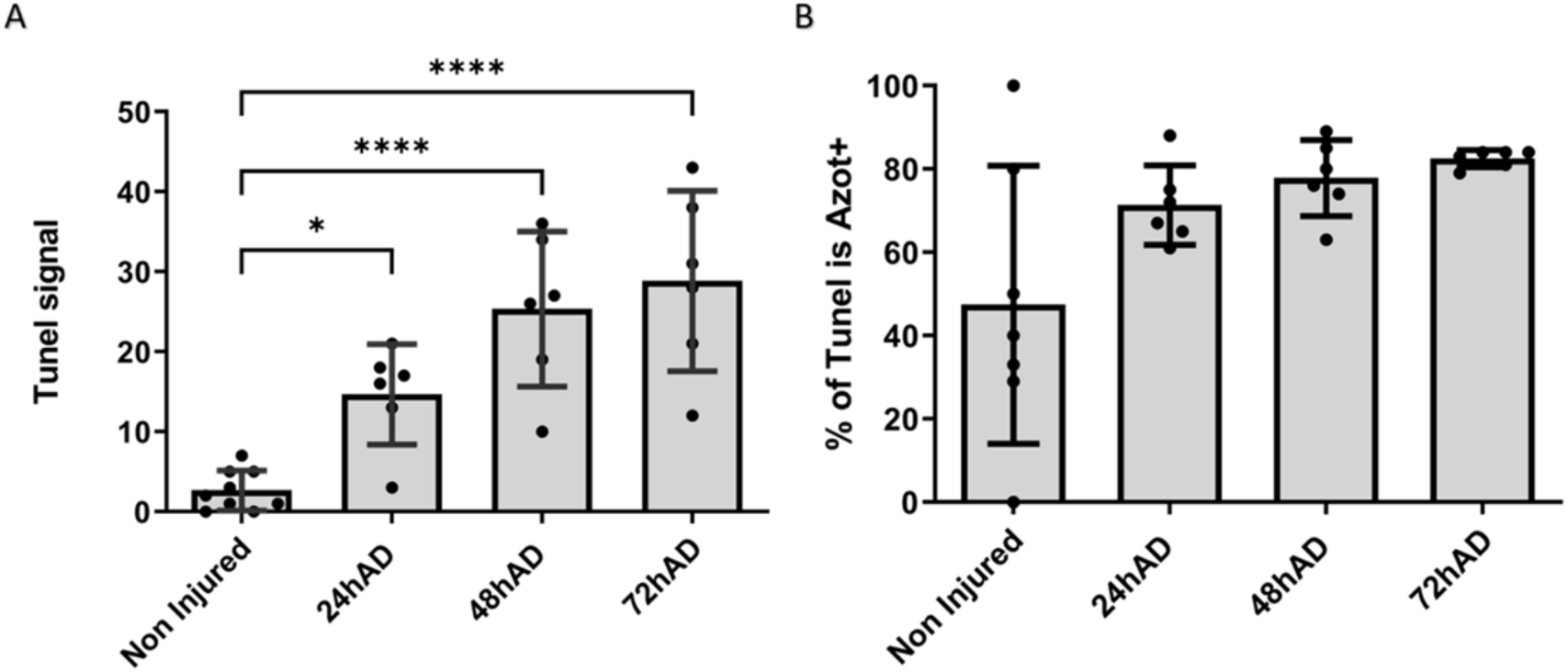
Apoptotic events occur in the brain after TBI and the majority of dying cells are loser. (A) Quantification of TUNEL signal after TBI. Statistical comparisons shown are only made with the “Non-injured” control. Error bars indicate SD; *P≤0.05; ****P≤0.0001. Statistical significance was calculated with one-way ANOVA and Dunnett’s post hoc test. (B) Percentage of TUNEL signal that colocalizes with Azot expressing cells. Error bars indicate SD. Genetic background: *ywf*; *azot*{KO-KI *azot::mCherry*} (Hz); +

Finally, we evaluated the percentage of dying cells, TUNEL positive, that colocalized with Azot positive cells (losers) (Figure 2B). In a non-injured scenario, this percentage is 47.4%. At 24, 48, and 72 hours AD, this percentage increased to a mean of 71.3%, 77.8%, and 82.5%, respectively, with less variability than the control. Collectively, these findings demonstrate that Azot is expressed in the optic lobe after TBI and that these loser cells are being eliminated through apoptosis.

### Loser cells after TBI are predominantly neurons and glia

We then wanted to investigate the identity of the loser cells in the TBI model and performed a double immunostaining for Repo (pan-glial marker) and Elav (pan-neuronal marker) at 48 hours AD (Figure 3A). Our results reveal that while 50% of the loser cells are Elav-positive neurons, surprisingly, 30% co-localized with the glial marker Repo, and 20% were negative for both markers (Figure 3B). These findings indicate that multiple cell populations in the brain, are susceptible to cell competition following traumatic brain injury.

**Figure 3.**
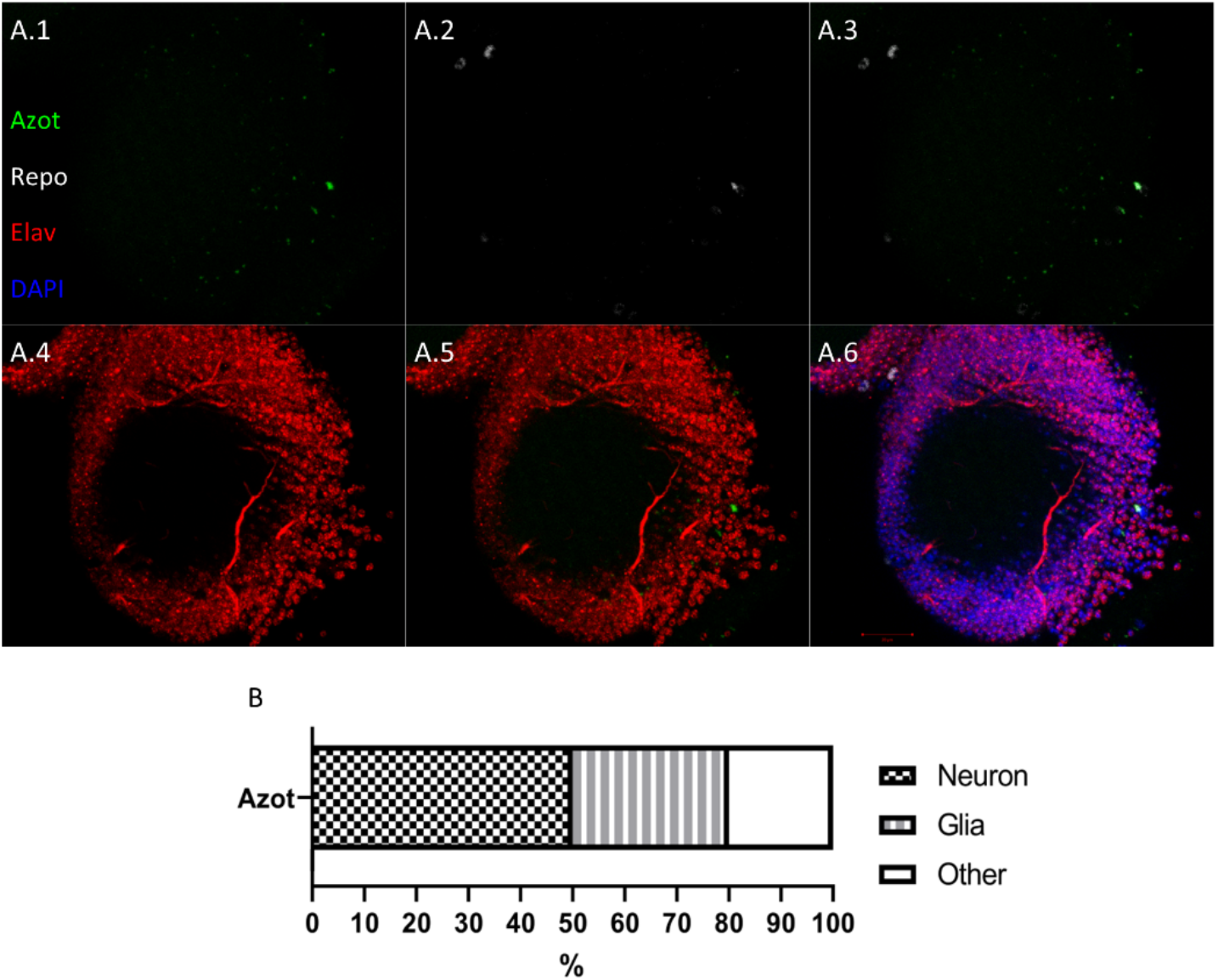
Loser cells upon TBI are predominantly neurons and glia. (A) Inset of a ROL 48h after TBI. Green channel is Azot (GFP), 647nm (here in white) is α-Repo staining, red channel is α-Elav, and blue channel is DAPI. A.1 is the green channel, A.2 is the Repo, A.3 is the merge of green and Repo, A.4 is red channel, A.5 is the merge of red and green channels, and A.6 is the merge of all the channels and DAPI. 8µm Maximum intensity projection. Scale bar is 20µm. (B) Quantification of Repo and Elav Azot-positive cells after TBI. 11 OL were analyzed. Genetic background: *ywf*; *azot*{KO-LexA}/+; 26xLexAop-*CD8::GFP*/+

### Loser cells accumulate in the absence of *azot* in the TBI model

It has been established that loser cells accumulate in the brain during aging when *azot* is absent^8^. Additionally, recent findings indicate that under conditions of *azot* depletion, loser cells express anti-apoptotic markers, suggesting that *azot* is not essential for loser cell elimination^10^. This points to the existence of redundant mechanisms that eventually lead to the elimination of these suboptimal cells -albeit with slower kinetics-, explaining their temporal accumulation. We investigated whether a similar accumulation of loser cells occurs in the brain following TBI when the *azot* gene is absent.

We utilized a homozygous (Hz) stock with a LexA insertion replacing the *azot* gene in its native locus, which was coupled with LexAop-GFP to label cells that would normally express *azot*^*31*^ (Figure 4A). Our findings indicate that in the absence of *azot*, cells attempting to express it accumulate over time (Figure 4B). Even in non-injured conditions, we observed an average of 22 cells per OL (compared to 9.5 cells when Azot is present). Following TBI, we found a continuous increase in these cells from 24 to 72 hours after damage. At 24 hours AD, we observed an average of 129 cells per OL, increasing to 165 cells at 48 hours AD, and 178 cells at 72 hours AD, representing percentage increases of 27.9% and 7.9%, respectively (from 24 hours AD to 48 hours AD and 48 hours AD to 72 hours AD). Interestingly, the number of Azot-expressing cells at 7 days after injury decreased by 37.64% compared to levels at 72 hours, with an average of 111 cells per OL. Since the absence of *azot* prevents cell elimination via cell competition, this reduction from 72 hours to 7 days likely indicates an alternative elimination mechanism, although it could potentially suggest that some cells eventually cease Azot expression and persist in the tissue. These results underscore the importance of Azot in regulating the damage response and point to additional processes involved in the clearance of suboptimal cells.

**Figure 4.**
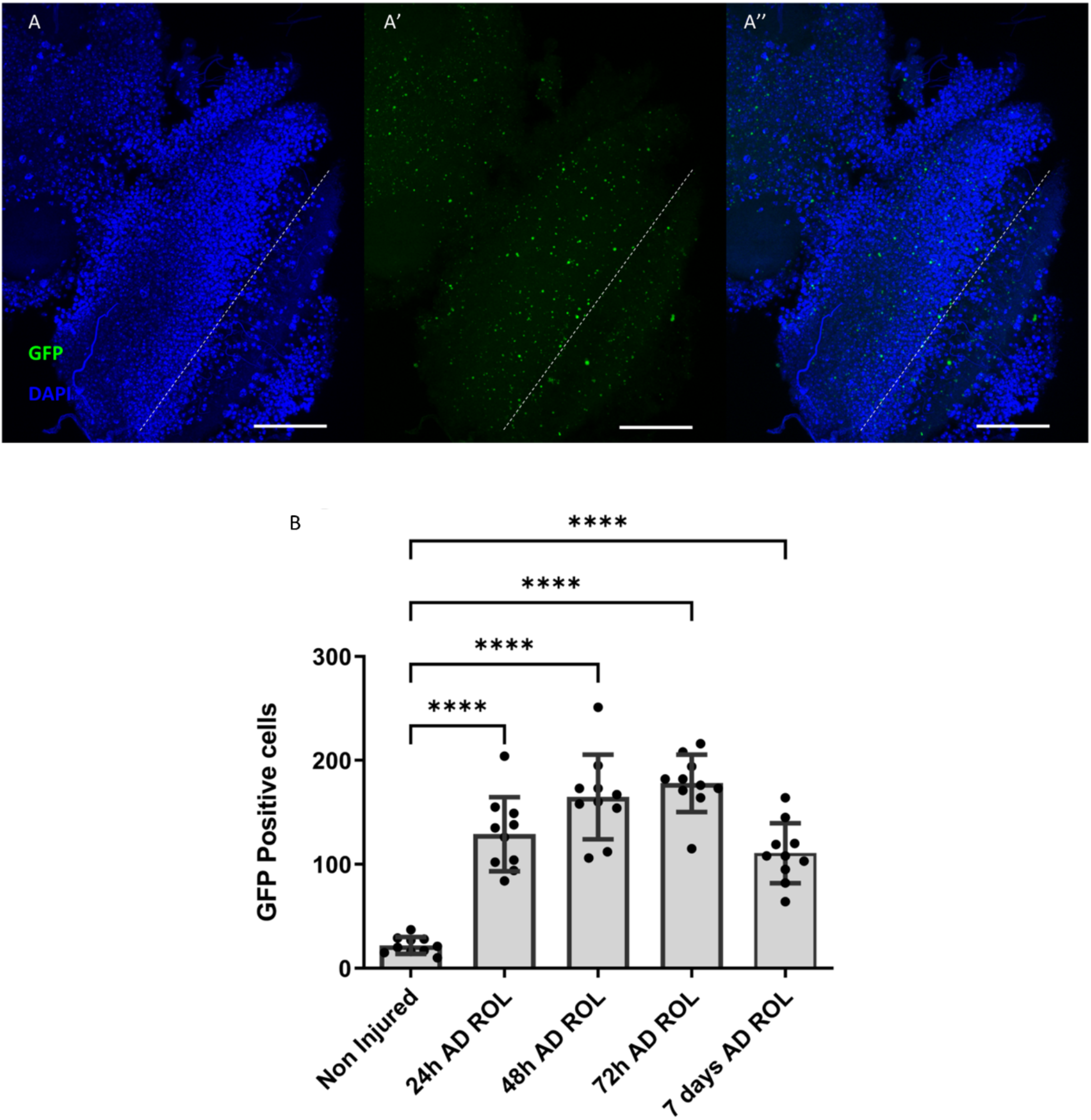
Loser cells accumulate in the OL in the absence of Azot. (A) Inset of a ROL 48h after TBI. A is the blue channel for DAPI, A’ is the green channel for GFP, and A” is the merge of blue and green channels. The dashed line indicates the wounded area. 40µm Maximum intensity projection. Scale bar: 50µm. (B) Quantification of the number of GFP-expressing cells in the OL. Statistical comparisons shown are only made with the “Non-injured” control. Error bars indicate SD; ****P≤0.0001. Statistical significance was calculated with one-way ANOVA and Dunnett’s post hoc test. Genetic background: *ywf*; *azot*{KO-LexA} (Hz); 26xLexAop-*CD8::GFP* (Hz)

## Discussion

Building upon the established link between TBI and cell competition and the downstream expression of Azot following Flower LoseB upregulation^26,27^, we characterized the temporal dynamics of Azot expression post-TBI. Our findings reveal a distinct expression pattern: a peak at 24 hours followed by a decline up to the 7th day AD. TUNEL assay confirmed that these Azot-expressing loser cells undergo apoptotic elimination. Interestingly, the contribution of cell competition to cell death increases from approximately 50% in non-damaged conditions to 80% in the damaged brain. The variability in this percentage progressively decreases from 24 to 72 hours AD, accompanied by enhanced colocalization of dead cells with Azot. These findings suggest that cell competition becomes the predominant elimination mechanism during the acute post-injury phase, in contrast with homeostatic basal conditions of the non-damaged control, where cell competition may or may not be involved in the cell death.

Interestingly, combined analysis of Azot-expressing cells and TUNEL-positive cells does not reveal a distinct “main wave” of cell elimination. Rather, the data suggests a progressive reduction in the timeframe of cell elimination, indicating that Azot-expressing cells undergo accelerated demise the further along from the initial damage. This enhanced cell competition may be attributed to the increased number of cells expressing Flower LoseB in the tissue, together with potentially unknown mechanisms that progressively shorten the period between Azot activation and apoptosis. This compressed timeframe likely prevents detection of Azot levels at later timepoints prior to caspase activation initiation. The post-TBI microenvironment^32^ likely potentiates this effect through multiple concurrent processes— neuroinflammation, necrotic signaling, immune cell infiltration, axonal degeneration, oxidative stress, and lipid peroxidation^23,28^—collectively reducing the average fitness in the wound periphery. The higher proportion of unfit cells likely intensifies intercellular fitness comparison, elevates Flower LoseB expression, and consequently increases and enhances Azot activation, thereby accelerating the identification and elimination of suboptimal cells from the injured tissue.

Recently, our group discovered that some neurons eliminated during Alzheimer’s disease are selected via Cell Competition^7,33^. Importantly, although controversial, enhancing the elimination of less fit neurons appears to improve motor behavior in affected flies. This represents a case of “beneficial death”; even though losing neurons is typically undesirable, in Alzheimer’s Disease, eliminating malfunctioning neurons may be preferable to having them persist and disrupt central nervous system function. Removing these dysfunctional neurons potentially facilitates neuroregeneration and/or the establishment of new synapses, while blocking their elimination worsens motor symptoms and decreases fly lifespan. These findings from neurodegeneration models may have parallel implications for TBI recovery, where the selective elimination of compromised cells could similarly benefit overall tissue function and recovery trajectories.

Intriguingly, in *azot* knockout conditions, the population of cells attempting to express Azot diminishes by the seventh-day post-injury. This finding suggests several possibilities: (1) activation of alternative elimination pathways with delayed kinetics^10^, (2) resolution of the “unfit” status in surviving cells, or (3) clearance through non-cell-autonomous mechanisms such as immune-mediated phagocytosis^10^. This observation highlights the complex and potentially redundant nature of quality control mechanisms in the injured brain. Further characterization of cell death in *azot*-deficient conditions through TUNEL assays would provide valuable insights into these compensatory elimination mechanisms in the TBI context.

Previous work from our laboratory in a *Drosophila melanogaster* Alzheimer’s Disease model demonstrated that cells in the brain positive for the loser fitness marker Flower LoseB were exclusively neurons, as they were negative for the pan-glial marker Repo and positive for the pan-neuronal marker Elav^7^. We found that in the TBI model, cell competition in the brain extends beyond neuronal populations to include glial cells. The cellular and structural complexity of the brain introduces several intriguing possibilities: competition may occur between distinct cell types (neuron-glia competition), glia may modulate neuronal competition without directly participating, or parallel competition systems may operate independently within neuronal and glial populations. Delineating these possibilities represents an important direction for future research.

This work advances our understanding of post-TBI cellular dynamics by characterizing the expression pattern of Azot and demonstrating its role in loser cell elimination across multiple cell types. These findings have significant implications for understanding brain injury response and repair mechanisms. Recent literature has begun to elucidate the complex interactions between neurons and glia in activating quiescent neural progenitors following injury^34^. Our future research will investigate how cell competition and the selective elimination of suboptimal cells in the injured area influence the regenerative capacity of the injured brain. By understanding these endogenous quality control mechanisms, we may identify novel therapeutic targets to enhance functional recovery following traumatic brain injury.

## Experimental Procedures

### Drosophila stocks

Stocks and crosses were kept at 25ºC, with a humidity level of 70%, in Vienna standard media with extra yeast. The following stocks were used: *azot*{KO; KI-*LexA::p65*}^10^, 26xLexAop-CD8::GFP (Bloomington Drosophila Stock Center, stock #32207), and *azot*{KO; KI-*azot::mCherry*} (this work).

### Azot knockin generation

The *azot* knockout founder line was described by Merino et al. 2015^8^. For the generation of the *azot*{KO; KI-*azot::mCherry*}, the cDNA of *azot::mCherry* was generated and inserted into a vector w+, AmpR, and the knockin was generated as described in Huang et al. 2009^35^, following the same strategy as the *azot*{KO; *Gal4*} described by Merino et al. 2015^8^. Primer sequences are available upon request.

### Traumatic Brain Injury

Before performing the procedure, the flies were anesthetized with CO_2_. Adult flies 2 days old were stabbed in the Right Optic Lobe (ROL) through the cuticle on the dorsal part of the head, as described in Moreno et al., 2015^26^ and Fernández-Hernández et al., 2013^25^, next to the pair of bristles that surround the retina. The filament was sterilized in 100% Ethanol each time a TBI was performed. Upon injury, flies were kept at 25°C until dissection.

### Brain Dissection and Immunostaining

Adult *Drosophila* brains were dissected in chilled S2 medium (SCHNEIDER’S DROSOPHILA MEDIUM), fixed overnight at 4°C in PFA 4%, washed in PBS-TritonX-1% (PBT) and mounted or prepared for immunohistochemistry. For immunohistochemistry, samples were incubated overnight in the primary antibody at 4°C, washed 3x 15’ in PBT, and incubated overnight in the secondary antibody at 4°C. Before mounting, the brains were washed 3x 15’ in PBT. The mounting was performed in “bridge” (with a spacer to avoid compressing the brains with the coverslip) as described in Fernández-Hernández et al., 2013^25^, using Vectashield with DAPI as mounting media. The following antibodies were used: rat anti-ELAV (DSHB Cat#7E8A10; RRID: AB_528218) and mouse anti-Repo (DSHB Cat# 8D12; RRID: AB_528448).

### TUNEL Staining

TUNEL (Roche) staining was performed according to the supplier’s protocol.

### Image Acquisition

Images were acquired using the confocal Zeiss LSM 880 Airyscan with the objective Plan-Apochromat 40x/1.4 Oil DIC M27. Any parameters, such as Laser Power or detector Gain, were set according to the state of the system at the specific moment of the acquisition. All OLs were imaged from the anterior to the posterior side. The depth is 40µm and the step between frames is 1µm.

### Quantification and Statistical Analysis

To quantify the number of cells, we used a macro in Fiji (ImageJ), available upon request.

Data from all samples were assessed for normality distribution using the Shapiro-Wilk test. Given that all the datasets followed a normal distribution, we employed ANOVA and proceeded with multiple comparisons using the Dunnett test to ascertain significant disparities between each sample/condition.

## Abbreviations

AD: After-damage
Dpp: Decapentaplegic
Hz: Homozygous
JNK: c-Jun N-terminal kinase
OL: Optic Lobe
PBT: PBS-TritonX-1%
ROL: Right Optic Lobe
TBI: Traumatic Brain Injury

## Author Contributions

A.G.G. and E.M. designed the experiments. A.G.G. performed the experiments. M.M.R. helped with the dissections and immunostainings. B.H. developed the *azot* knockin. A.G.G. and M.M.R. wrote the manuscript.

## Acknowledgments

We thank Bloomington Stock Center for flies; the technicians at the Champalimaud Fly Platform for support with stock maintenance; and the ABBE platform for microscopy support. M.M.R. was supported by an FCT - Fundação para a Ciência e a Tecnologia – PhD studentship (SFRH/BD/138537/2018). Portuguese national funds supported this study through FCT in the context of the project UIDB/04443/2020 and the European Research Council (Consolidator Grant to E.M.: ‘‘Active Mechanisms of Cell Selection: From Cell Competition to Cell Fitness’’). Fly platform was supported by CONGENTO LISBOA-01-0145-FEDER-022170, co-financed by FCT (Portugal) and Lisboa2020, under the PORTUGAL2020 agreement (European Regional Development Fund). The Portuguese Platform of Bioimaging funded ABBE platform - LISBOA-01-0145-FEDER-022122.

